# Disentangling the shark and ray trade in Indonesia to reconcile conservation with food security

**DOI:** 10.1101/2020.12.08.416214

**Authors:** Andhika P. Prasetyo, Allan D. McDevitt, Joanna M. Murray, Jon Barry, Firdaus Agung, Efin Muttaqin, Stefano Mariani

## Abstract

Indonesian marine resources are among the richest on the planet, sustaining highly diverse fisheries and includes the largest elasmobranch landings in the world, making Indonesia one of the world’s largest exporters of shark and ray products. Socio-economic and food security considerations pertaining to Indonesian communities add further layers of complexity to the management and conservation of these vulnerable species. This study investigates how shark and ray trade flows in and out of Indonesia and attempts to examine patterns and drivers of the current scenario. We identify substantial discrepancies between reported landings and declared exports, and between Indonesian exports in shark fin and meat products and the corresponding figures reported by importing countries. These mismatches are estimated to amount to over $43.6M and $20.9M for fins and meat, respectively, for the period between 2012 and 2018. Although the declared exports are likely to be an underestimation because of significant unreported or illegal trading activities, we find that domestic consumption of shark and ray products plays a significant role in explaining these discrepancies due to the increasing local demand for meat. The study also unearths a general scenario of unsystematic data collection and lack of granularity of product terminology, which is inadequate to meet the challenges of over-exploitation, illegal trade and food security in Indonesia. We discuss how to improve data transparency to support trade regulations and governance actions, by improving inspection measures, and conserving elasmobranch populations without neglecting the socio-economic dimension of this complex system.

## 1. Introduction

The rapid depletion of sharks and rays (hereafter referred to as just ‘sharks’) in many marine ecosystems is now recognized globally [1, 2]. Conservative life-histories [3] make sharks vulnerable to fisheries overexploitation [4, 5], which in turn can destabilise ecosystem structure [6] and ultimately decrease global functional diversity [7]. High market demand for high value shark products is the main reason for the overexploitation of shark resources [1, 8], which ultimately leads to illegal trade [9]. Much of this trade involves poorly reported catches from Eastern and Western Pacific countries, which supply, for instance, global shark fin markets [10, 11]. Understanding and regulating such trade is challenging because shark products are extremely diverse in both their usage and their value, and are processed in a myriad of different ways (Figure 1) [12-14].

**Figure 1.**
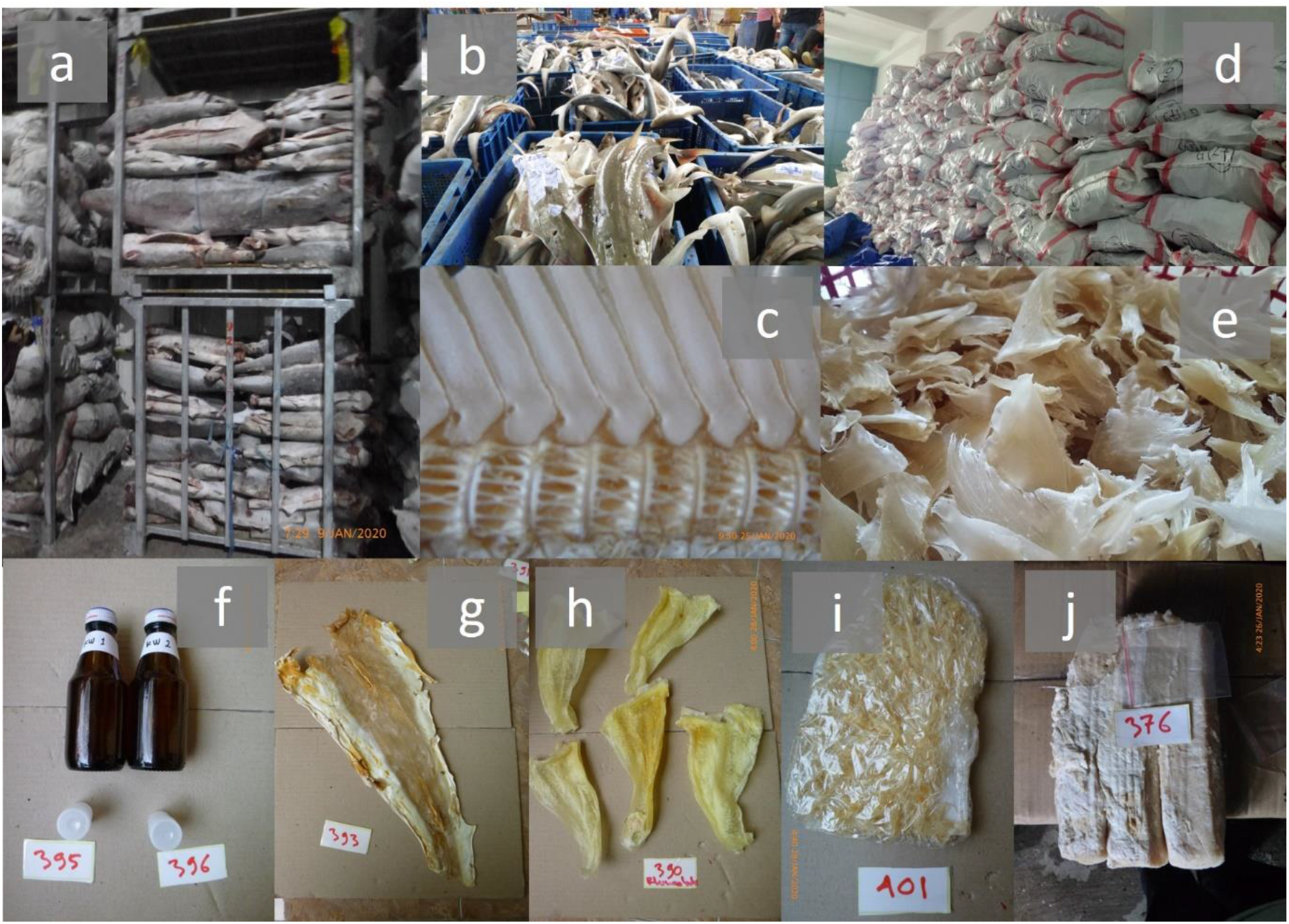
Storage, appearance and diversity (export commodities) of shark products: frozen shark trunks in cold storage (a), fresh rays landed in Indramayu, (c) ray cartilage, (d) stock pile of controlled species waiting for quota, (e) peeled shark fins, (f) shark oil, (g) peeled shark skin, (h) peeled ray fins, (i) “hissit” noodle-like from shark fins, and (j) shark salted meat.

A few regions of the world represent remarkable hotspots for elasmobranch diversity, making them focal targets for biodiversity conservation. Indonesia, with its many islands and diverse habitats at the interface between two ocean basins, is one such region, believed to harbour about 20% of global shark diversity (119 of 509 living sharks; 106 of 633 living rays), covering the whole spectrum of functional traits, from highly migratory oceanic species, to reef-associated, and sedentary bottom-dwelling coastal endemic taxa [15-17]. Indonesia is also the fourth most populous country in the world, with many communities traditionally associated with the sea [18]. This makes elasmobranch conservation management in Indonesia problematic, due to diverse and unregulated small-scale fisheries, high incidences of illegal fishing, and unsystematic data collection. Moreover, Booth, Muttaqin, Simeon, Ichsan, Siregar, Yulianto and Kassem [19] report that 86% of all Indonesian fisheries surveyed catch sharks incidentally or as by-catch and whole fishing communities also exist that target sharks exclusively, and in some cases even certain species in particular, using tailored gear [19, 20]. Between 2007-2017, Indonesia was the largest reported contributor to global shark landings, with a mean catch of 110,737 mt per year [21, 22]. The paired trends of depletion and exploitation – in such a biodiverse context – call for global attention to identify effective mechanisms to ensure sustainability of shark resources. This includes improving reporting, introducing regulations and ensuring compliance (e.g. through the Convention on International Trade in Endangered Species of Wild Fauna and Flora (CITES) framework [23] and other approaches [24], with the ultimate goal of identifying a balance between preserving wildlife and sustainable resource use.

Globally, market demand of shark products is stable, especially fin products [21]. However, since 2015, a dramatic increase was observed in the export of meat products in Indonesia. This has been linked to emerging trammel net by-catch, as a consequence of the ban on shrimp trawling [25]. Much of these landings are believed to include vulnerable/endangered species, including several currently listed in the regulatory trade annexes of CITES. Since sharks are processed in many ways, this poses challenges to CITES requirements (i.e. legality, sustainability, and traceability) and other regulatory frameworks [26]. The large amount of caught biomass, over a vast and diverse coastline, and the limited facilities and resources for inspection also add obstacles to effective monitoring of shark trade in Indonesia.

Shark conservation remains a high priority topic in marine ecology, but in many circles the focus is almost entirely on the goal of species conservation, with little emphasis on socio-economic aspects and limited evaluation of the trade-offs among the different stakeholders [27, 28]. This study aims to reconstruct the current state of shark trade in Indonesia in order to lay the foundations for a remodelled management framework for the world’s most diverse and vulnerable marine vertebrate resources. To do so, we: i) collate and summarise data on landing trends, ii) investigate domestic trade flows, iii) examine import/export discrepancies, iv) identify factors, challenges and solutions to maximise ecological and socio-economic benefits.

## 2. Material and methods

National shark production statistics were compiled from 1950 to 2017, taking into consideration that fisheries data collection started improving gradually from 2005. These considerations included accounting for the introduction of the CITES regulations (which started requiring the official recording of nine and seven categories of sharks and rays, respectively). In 2017, there was a significant change in national data collection operations, which included marine and fisheries sectors, which introduced the so-called “one-data” policy. This policy is designed to provide a regulatory framework and standard mechanisms to the principles of data interoperability among stakeholders [29-31]. Those official statistics were combined with the global capture production database from the UN Food & Agriculture Organisation [22] to provide better insight of both national and international sharks trade in Indonesia. Moreover, we defined ‘controlled species’ as all sharks and rays that are listed in CITES’ annexes. Trade activities that fail to comply with national or international laws for such ‘controlled species’ are deemed ‘illegal trade’.

The domestic trade flow was examined by mining datasets from 46 fish quarantine offices across Indonesia, which included information about location of sources and destination, type of products, volume and estimated value [32]. The volume of domestic shark product exchange between source and destination locations was then plotted using the R package “network3D” [33]. To improve clarity, domestic trade was filtered to flows larger than 10 ton.

Shark trade data to examine exports reported by Indonesia (hereafter ‘Export from Indonesia’) and imports from Indonesia that were reported by other countries (hereafter ‘Import from Indonesia’) were derived from the FAO Fisheries Statistics [34] and the Agency for Fish Quarantine and Quality Insurance [32] for a seven-year period (2012–2018). This analysis period was selected because the FAO Fishery Commodities and Trade statistical collection [34] included shark import and export records only starting from 2012. Data were then filtered by selecting i) type of trade flow (export, import or re-export), ii) source or destination country, and iii) harmonized system (HS) code (a code that consists of an internationally standardized system of numbers to classify traded products and commodities). Given the fluctuations in export and import value of fin and meat products, we estimated trade record mismatches by averaging the values between exports and imports over the whole 2012-2018. Bilateral trade flows between Indonesia and importing countries were represented using Circos [35]. Calculation and visualisation were performed in R 3.6.1 [36]. Discrepancy between Indonesia and bilateral trade partners were traced using the method detailed by Cawthorn & Mariani [37] by subtracting the export figure reported by Indonesia from the corresponding volume reported by each partner country. The results were aggregated for the study period and for examined commodities, unless otherwise specified. Additional information about data sources can be found in Supplementary Table S1.

## 3. Results

### 3.1. Production statistics

Indonesia ranks as the world’s top shark landing country in terms of quantity, while its imports are negligible. According to government production statistics, annual elasmobranch production has rapidly increased between the 1970s and 2000, becoming relatively steady over the past decade (2005-2014), oscillating between approximately 90,000 to 120,000 tonnes per year, with a 10-year annual average of 107,623 (SD 12,932) tonnes [22, 29, 30]. Sharks generally amounted to just over half of landings, with the situation reversed in the last six years, when rays peaked to account for up to two thirds of reported catches in 2016 (Figure 2). It is also important to note that, although MMAF monitoring systems currently classify sawfishes as ‘sharks’, for the purpose of this study, we placed them among the rays, in line with their systematic classification (Batoidea: Rhinopristiformes) [17].

**Figure 2.**
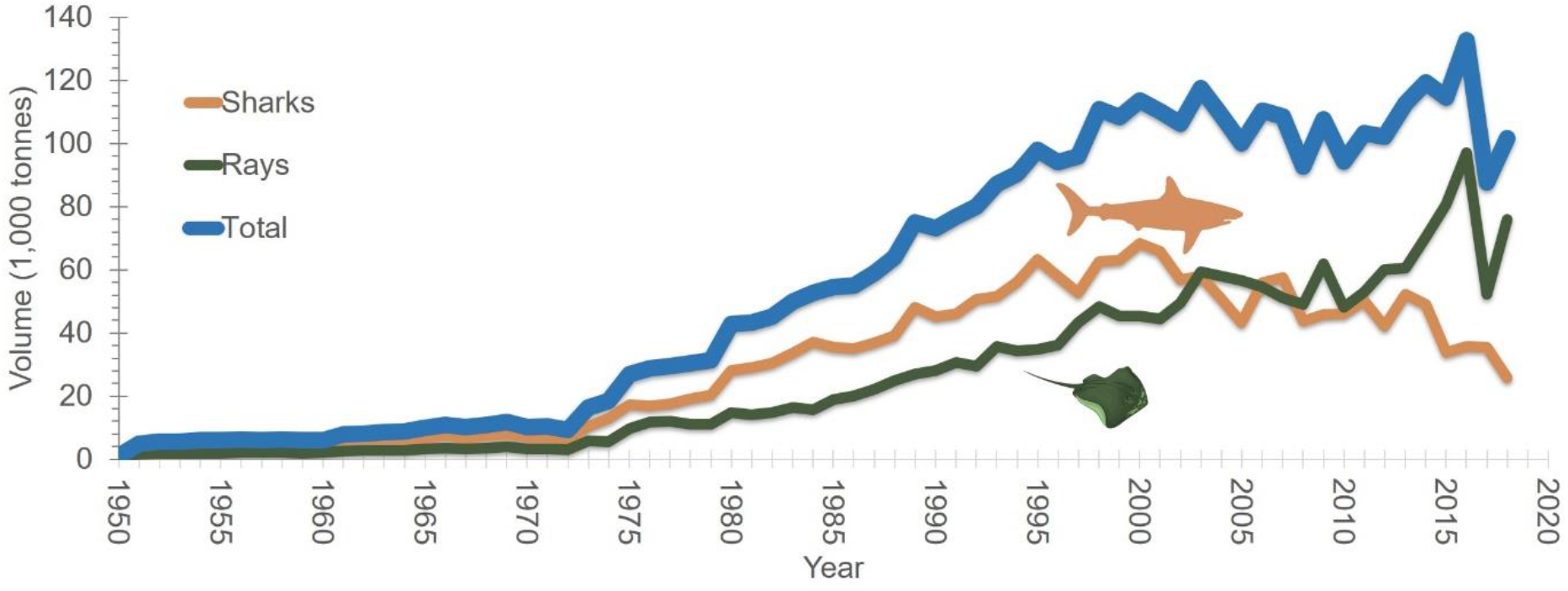
Shark landing in Indonesia 1950-2018. [22, 29, 30]

Indonesian shark production records are rather coarse in terms of taxonomic granularity, with limited species-specific information. Instead, landings are grouped into broad categories, as collected by MMAF, e.g. requiem sharks (other Carcharhinidae) and thresher sharks (Alopidae) which made up most of the shark production over the past 14 years, contributing 51% and 22%, respectively (Figure 3a). Shark production from 2005 to 2018 fluctuated for each species group, but generally declined since 2016. Requiem (Carcharhinidae) and mackerel (Lamnidae) sharks have shown stable volume over time. CITES-listed silky sharks (*Carcharhinus falciformis*) fall within the broader requiem shark group (other carcharhinidae), while tiger shark (*Galeocerdo cuvier*), oceanic whitetip shark (*C. longimanus*) and blue shark (*Prionace glauca*) were only recently put into separate categories in 2015.

**Figure 3.**
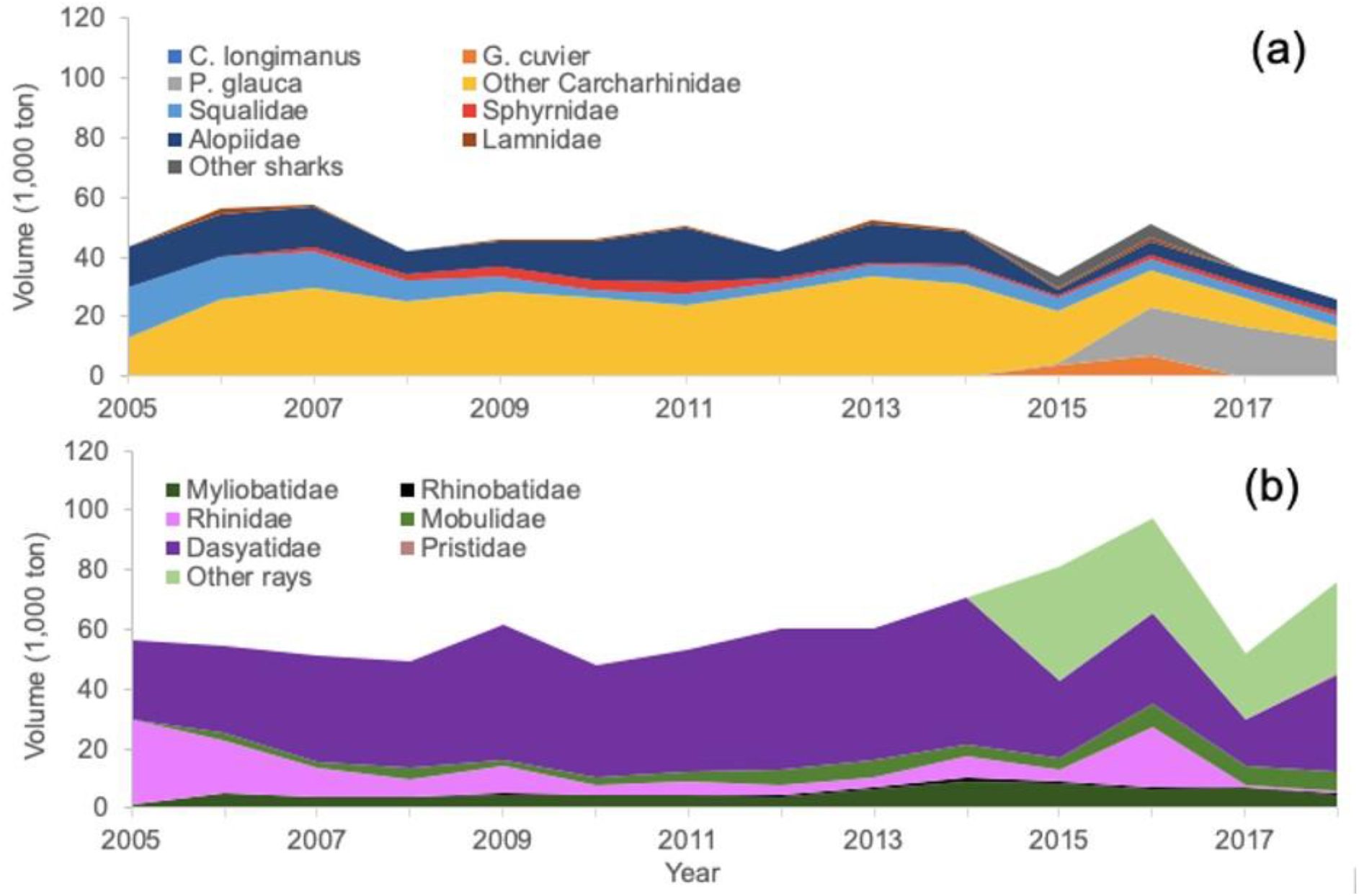
Sharks (a) and ray (b) landing and composition in Indonesia by species group 2005-2018 [22, 29, 30].

Stingrays (Dasyatidae) made up most of the ray production over the past ten years (56%), followed by wedgefishes (Rhinidae) (13%) and eagle rays (Myliobatidae) (8%). Ray production for most species has generally increased over time, although wedgefishes saw declines between 2005 and 2010 (Figure 3b). Much uncertainty is associated with catch records because various species and families are misidentified and mistaken, such as the case of ‘sawfishes’ (Pristidae) and ‘sawsharks’ (Pristiophoridae), or ‘wedgefishes’ (Rhinidae) and ‘guitarfishes’ (Rhinobatidae).

Indonesia has 11 Fisheries Management Area (FMA) that overlap with provincial jurisdiction’s areas (34 provinces). During the 2005-2018 period, nearly 1,488,006 ton sharks and rays were landed across Indonesia’s 11 FMAs. FMA 711 (North Natuna Sea) and FMA 712 (Java Sea) were the major contributors, with 387,685 and 324,331 ton, respectively (Figure 4). In these two major areas, ray landings were substantially greater than shark catches.

**Figure 4.**
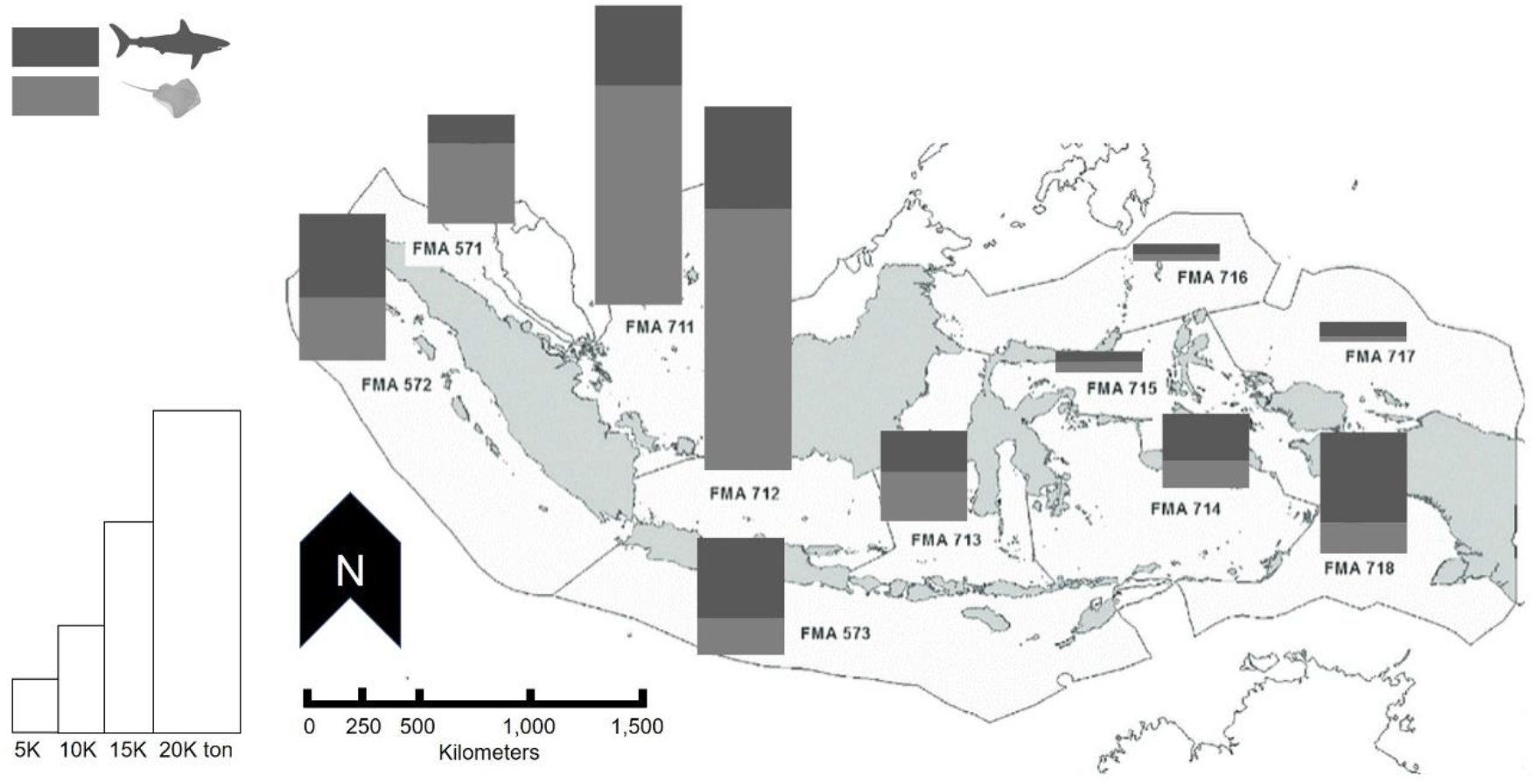
Cumulative volume of shark and ray landing by Fisheries Management Area (FMA) during 2005-2018 [22, 29, 30].

### 3.2. Domestic trade statistics

Based on national statistics, in 2018, the export of shark products was only just over 11.7% (11,867 ton) of landing data (101,707 ton), and only around 3.7% (24,839 ton) over the whole period between 2013 and 2018 (668,955 ton). We looked in more detail at the two main factors that can explain these figures: the substantial role of domestic consumption, and the potential for unreported/inaccurate trade figures.

Regarding domestic use, insights can be gleaned from documented internal trade flows. As a large archipelagic country, even the internal supply chain is complex: involving several actors and transit locations. There are several main source provinces of shark commodities, such as Bali, Papua, West Papua, East Kalimantan and Bangka-Belitung Provinces (Figure 5a), with Bali and Papua together accounting for 68.2% of the outflow at 10,587 ton. The Bali province also plays a role as a transit hub prior to subsequent shipping to Jakarta and East Java Provinces (Surabaya) (Figure 5b), which are the two main international export hubs. In such a complex scenario, and with the current uncertainties associated with product names, it is easy to imagine that some controlled species, while being allowed to be traded domestically, can become virtually untraceable after a few steps along the supply chain. It is worth noting that Aceh and Medan (North Sumatra) have no record of any trade, which is surprising, given their geographic proximity to key international markets, such as Singapore and Malaysia. Additional information about domestic flow can be found in Supplementary Figure S2.

**Figure 5.**
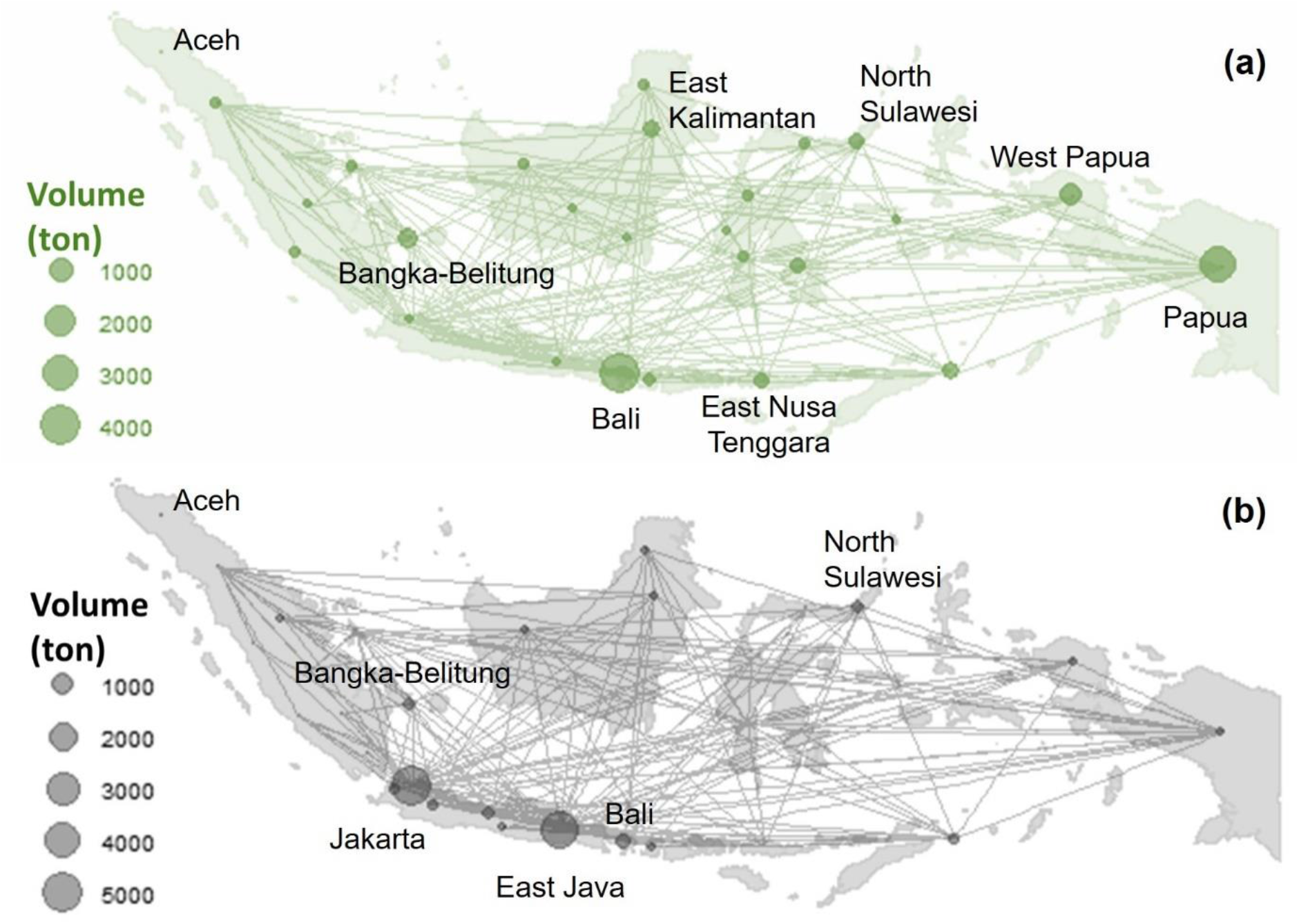
Domestic trade network of fin and meat products across Indonesia region within 2014-2018 (ton) by source (a) and destination provinces (b); provinces with label indicate significant contribution. [32]

### 3.3. International trade statistics

Shark meat is mainly for domestic consumption, while other valuable shark parts, such as fin, cartilage and other derivatives are export commodities. Between 2013 and 2018, exported shark products increased steadily and reached a peak in 2017 of 8,320 ton (Figure 6a). Over 70% of the exported product is still dominated by meat, except in 2016, where export of fins (878 ton out of 3,002) and cartilages (1,346 ton out of 3,002) was substantial (respectively 29% and 45% of the total). Indonesia also imported shark products, mainly the small-sized fins that are processed into *hissit* (shredded fins; noddle-like). However, the volume is negligible, amounting to just 155 ton throughout the 2012-2018 period. Products from the two main export hubs (Jakarta and Surabaya) mainly were shipped to Japan, Singapore and perhaps even China and Hong Kong. In recent years, export of live sharks has also increased steadily, almost doubling every year (Figure 6 Figure 6b) and are likely associated with recreational purposes. This demand targeted the coral reef associated species, such as black-tip reef shark (*Carcharhinus melanopterus*), zebra shark (*Stegostoma fasciatum*), bowmouth guitarfish (*Rhina ancylostoma*) and whitespotted whipray (*Himantura gerrardi*). The living sharks are mainly exported to China, Hong Kong, USA and Malaysia.

**Figure 6.**
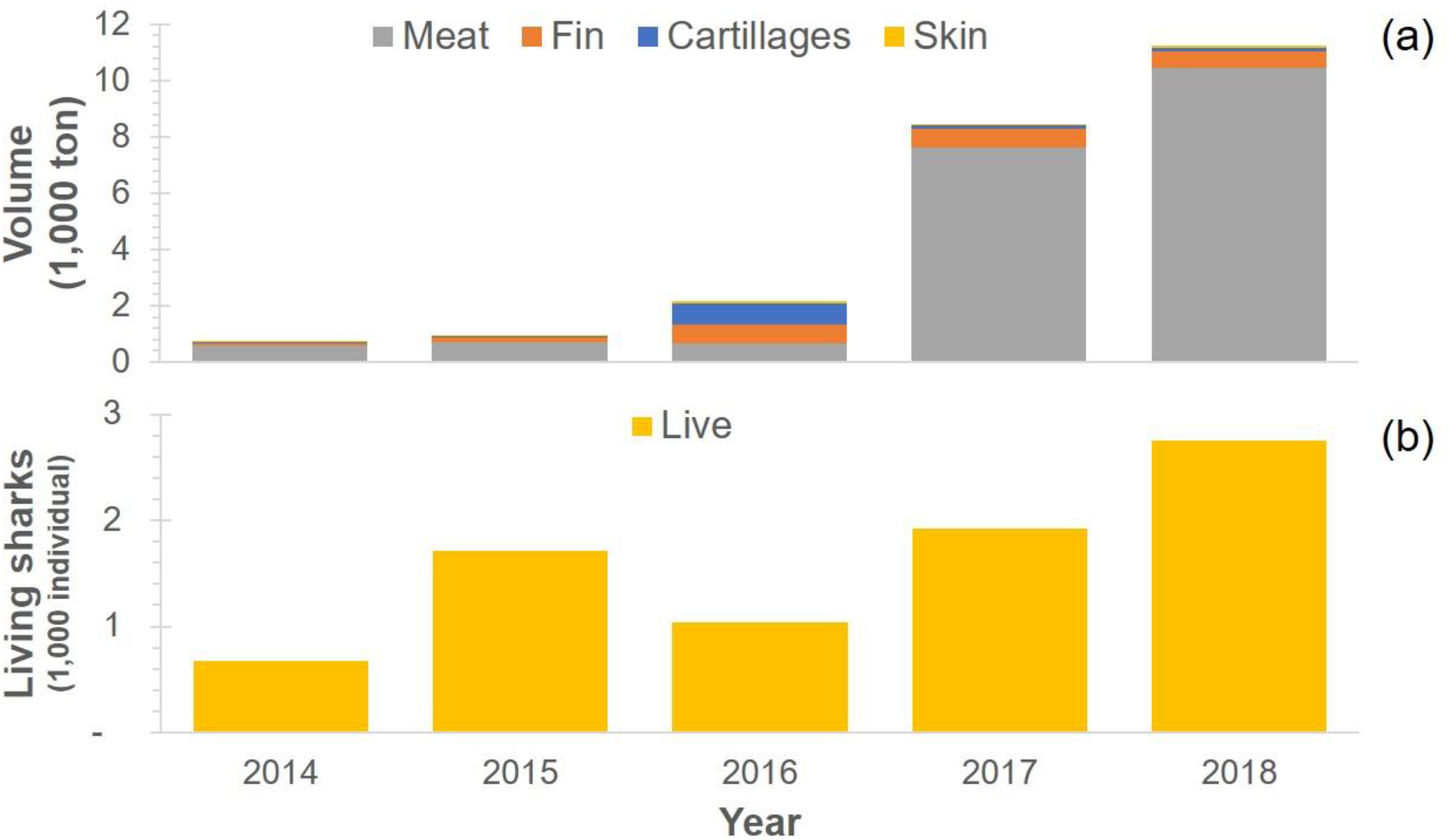
Export volume by products in 2014-2018 (a) and export for live sharks and rays in 2014-2018 (b). [32]

### 3.4. Data discrepancy

We extracted export-import data from FAO Trade Statistics on shark products, from 2012 to 2018, treating ‘fins’ and ‘meat’ separately. ‘Export’ was defined as the product figures reported by Indonesia as traded out to other countries (‘partners’), while ‘Import’ represented the amount of produce that each trading partner declared as being imported from Indonesia [22]. We found a substantial level of misreporting in the fin trade (Figure 7a). In some cases, Indonesia reported less than what the importing countries declared (e.g. Hong Kong reporting 440.5 ton more than what was stated by Indonesia), and in other instances it was the importing partner reporting less incoming trade from Indonesia (e.g. Singapore declaring 521 ton less than what was recorded by Indonesia). Similarly, this phenomenon was also revealed in the meat trade (Figure 7b), with the notable case of Malaysia, which reports nearly 9,000 ton more incoming trade than what was shown by the Indonesian export records. On average, the discrepancy of fin and meat products were 54.4% (1,462 ton) and 47.1% (13,138 ton) of the export volume reported by Indonesia (2,689 ton and 27,871 ton). This discrepancy was valued at 43.6 million US$ for fin and 21 million US$ for meat products. Additional information about discrepancy can be found in Supplementary Figure S3.

**Figure 7.**
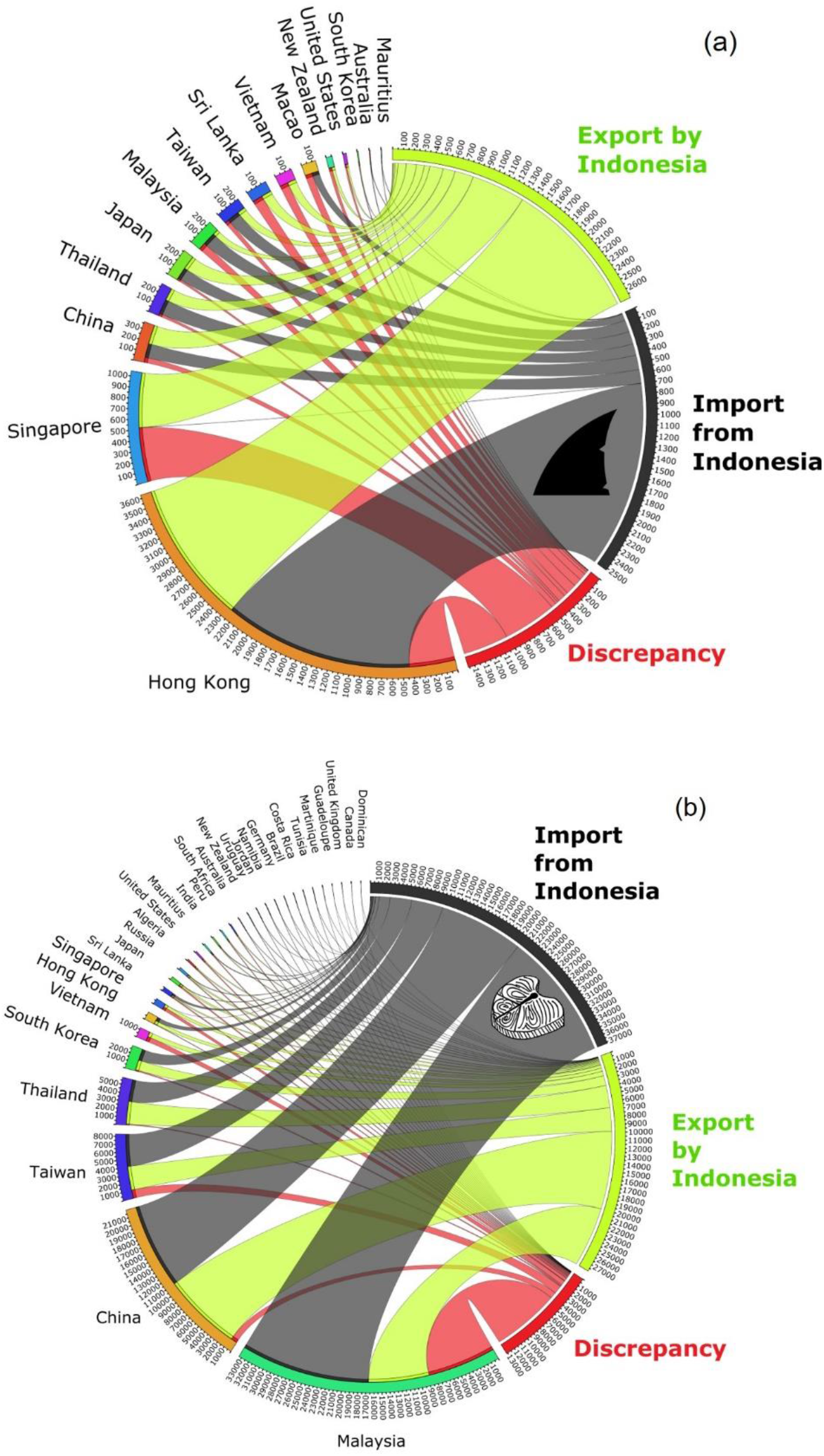
Trade flow and discrepancy of shark fin (a) and meat (b) products between Indonesia and its main trade partners, in tonnes, within the 2012-2018 period. Legend: Discrepancy (RED flow); the exported volume declared by Indonesia (GREEN flow), and the corresponding amount declared by each importing country (GREY flow). Source: [34]

**Figure 8.**
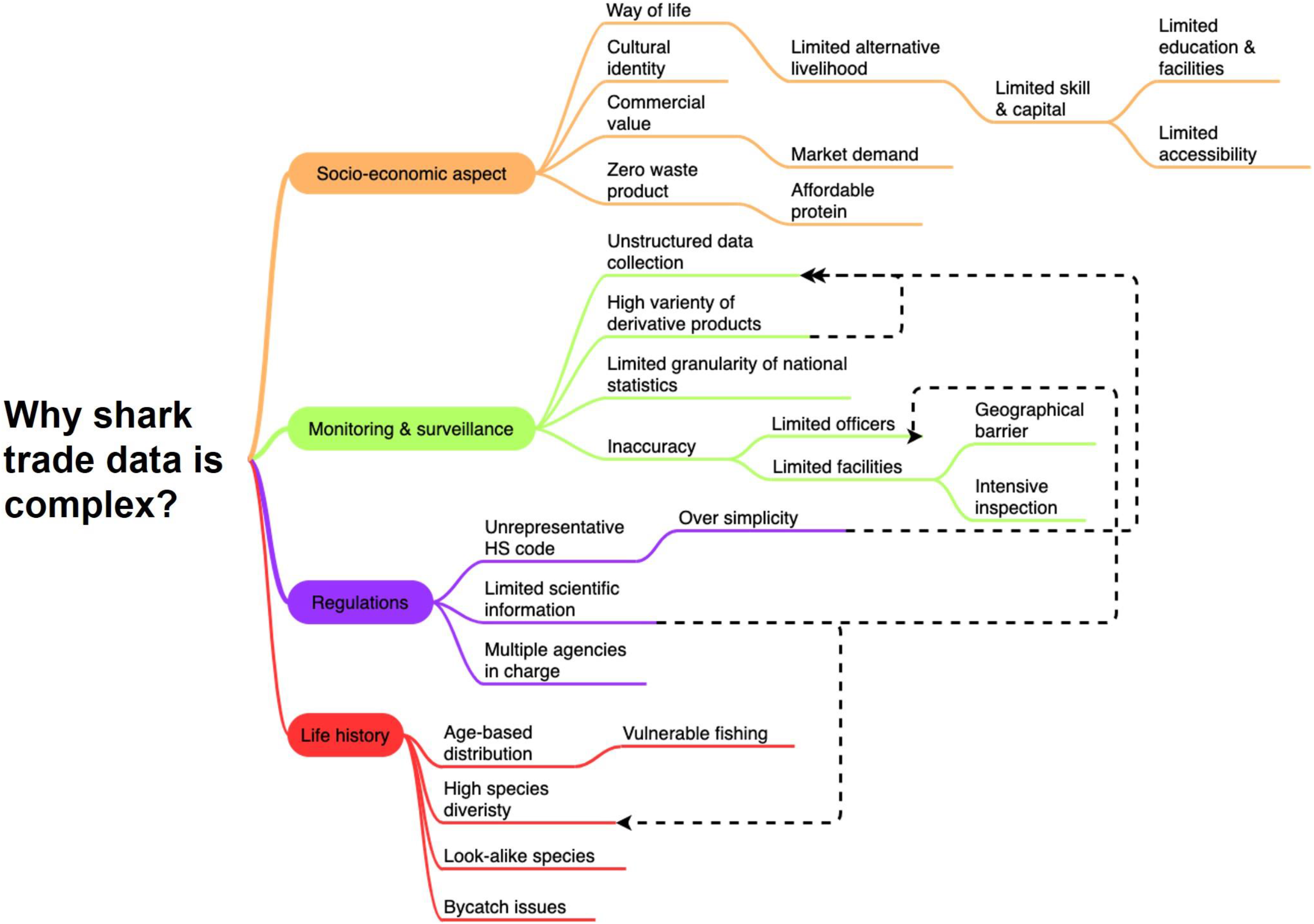
Causal diagram to explain the complexity of shark trade in Indonesia.

## 4. Discussion

Anthropogenic impacts on functional diversity of marine megafauna, their ripple effect on ecosystem structure [6, 38], and greater awareness of the value of marine predators when alive [39] has led to increased global attention to shark conservation. However, without a comprehensive understanding on the socio-economic dynamics around shark resources, such as the importance of shark fisheries for food security and the value chain associated with the international trade in shark products, conservation success is unlikely to be attained in the medium to long term [40, 41].

This study reveals significant inadequacies of existing trade statistics for the nation that lands the world’s largest volume of elasmobranchs. Those inadequacies reflected in four divergence issues, namely (1) volume gap between landing and export, (2) information gap between main landing site and main supplier at the domestic level, (3) volume gap between export and reported import by trade partners, and (4) impression gap between fisheries policy and bycatch reduction.

The mismatch between landing and export volumes, the inability to reliably partition landings into domestic consumption and international components [12], and the poor taxonomic granularity of catch (and trade) compositions, epitomise the grand challenges facing the Earth’s socio-ecological systems. This is especially important in highly populated, developing and biodiverse regions. Indeed, shark products sustain a diverse array of markets, from lucrative demands for traditional delicacies, supplies for medicines and cosmetics, curios, and substantial provision of food for local communities [12, 42]. The diversity and vulnerability of the living resources exploited, and the complex trade routes of their derivatives, calls for a step change in the ways data are recorded, fisheries are managed, and commercial activities regulated.

Global trends of trophic shifts following exploitation, where rays increase after shark removal [6] appeared to be mirrored into Indonesia landing record: within the last six years (2013-2018), ray landing volume (436,853 ton) was nearly twice as much as the figure for sharks (232,102 ton). As secondary catches, sharks have added value for fisheries and bycatch mitigation strategies remain inadequate to conserve these fragile creatures [2]. Some species of sharks have begun to be recorded separately to accommodate international trade measure, i.e. CITES. Requiem sharks (other Carcharhinidae) and thresher sharks (Alopidae) were the highest contributors to shark catches while ray was dominated by stingrays (Dasyatidae) and wedgefishes (Rhinidae). This is a major concern, as silky sharks (*Carcharhinus falciformis*), falling in the ‘other Carcharhinidae’ group, and wedgefishes, have both recently been added to international trade restriction. Moreover, the two main fishing management areas (FMA) that contributed the largest sharks catches (Java Sea and North Natuna Sea) are well known as fishing ground for wedgefishes and guitarfishes, and important bases for several fishing fleets that typically fish across other FMAs, such as FMA 713 (Makasar Strait) and FMA 718 (Arafura Sea).

Further issues are caused by discrepancies between landings and domestic trade flows in Indonesia. We found that Papua and Bali Provinces were the main market sources of shark products, but the main landing sites of shark catches were the North Natuna Sea (FMA 711) and Java Sea (FMA 712). A high domestic consumption in the main landing provinces which located in the densest population islands (Java and Sumatra) in Indonesia could be main factor explaining the mismatch. Domestic consumption aside, unsystematic data recording possibly contributes to confound the picture. Anecdotal information indicates that many sharks caught in the Arafura Sea (FMA 718) are shipped to Jakarta using cargo ship and landed in the cargo port, where they are recorded as ‘product’ instead of catches by the Fishing Port Authority in Jakarta. It was also noticed that the Aceh Province in Sumatra Island shows no domestic trade record, which suggests unreported exchanges among neighbouring provinces or even direct international trade with bordering countries, such as Malaysia and Singapore.

International trade over the most recent six years investigated (2012 – 2018), reveals a cumulative export of 2,689 tonnes of fins and 27,871 tonnes of shark meat reported by Indonesia, valued at approximately 41 and 40 million US$ respectively. Such products are mainly exported to Hong Kong, Malaysia, Singapore, China and Thailand. Hong Kong was the main market of fin products while Malaysia was the main destination of meat products (which mostly consisted of the fresh meat of rays). These bilateral trade depictions do not attempt to match shark commodities that were imported only to be subsequently exported (re-exports), as FAO data suggest that such re-exports are negligible.

Since there was a significant difference between the export and import value of shark products, the mismatch value was estimated using the average value between export and import in 2012-2018. Analysis of international trade shows significant discrepancy between export and import figures for fins and meat products by 1,462 ton and 13,138 ton respectively. This mismatch amounts to 54.4% of the total 2,689 ton export declared in the fin trade, which is valued at approximately 43.6 million US$ (based on the estimated value of 29,800 US$/ton). Gaps are mostly caused by fin trade with Singapore (under-reporting) and Hong Kong (over-reporting), by 521 and 440 ton respectively. On the other hand, there was a mismatch of 47.1% of the reported export in the meat trade, a value of approximately 21 million US$ (based on the estimated value of 1,600 US$/ton), most of which is due to the underreporting of products putatively imported by Malaysia (nearly 9,000 ton). This highlights the immense economic loss due to the mismatch in fin and meat products. These gaps could be filled, at least to some extent, by increasing granularity of shark product types in the World Customs Organization (WCO) Harmonised System (HS) codes. Currently shark products can be traded into 12 HS categories, which mostly emphasize differences in processing, yet invariably aggregate all ‘sharks’, ‘dogfish’, and ‘rays’ in the same group (Supplementary Table S4). This is of course insufficient to accommodate the high diversity of shark and ray species that regularly feature in traded products. It also reinforces concerns regarding the effectiveness of international measures to combat illegal trade [43, 44]. Similar findings on trade discrepancy between Hong Kong and its partner countries highlighted the importance of comprehensive data recording on shark fin trade [13]. It also advocates for the authorities to improve their capacity to reduce the risk that illegal products might contribute to such gaps.

The export of shark products (especially meat) increased dramatically since 2015, which could be connected with the introduction of a fishing moratorium by the Ministry for Marine Affairs and Fisheries (MMAF). MMAF issued decree no. 2/2015 concerning a trawl and seine-net ban in the Arafura Sea (FMA 718), in order to address shrimp stock depletion [45]. The subsequent shift from trawling and seine-netting to trammel-net activity led to a significant increase of shark bycatch. Within two years (2016-2018), processing plants in Jakarta have rapidly expanded shark product supply, leading to a “cobra effect” [46] whereby an attempted solution to a problem (i.e. overfishing of shrimp resources) actually makes the problem worse, and/or creates other unintended, problematic consequences (i.e. overfishing of endangered elasmobranchs). Current management should be reconsidered to attain a better trade-off of conservation and management measures [47]. Recently, an increased interest in displaying sharks alive in public aquaria and theme parks has led to higher demand of live sharks. China, Hong Kong, USA and Malaysia are the main market for such commodity, which usually comprise coral reef associated species. This emerging demand is anticipated to add complexity and additional challenges to trade regulations in relation to biodiversity protection and animal welfare.

There are now 48 species of Elasmobranchs listed in the CITES’s Appendices as of 2019. Of these, 30 are distributed in Indonesian and adjacent waters. Other authors have debated the effectiveness of the Convention’s measures [23, 48], but the Indonesian context is unique in its complexity, whereby high species diversity, high harvested biomass, complex internal trade routes, local population needs, and poor reporting and the potential for illegal wildlife trade all compound to set major challenges for the sustainable management of sharks and rays. Due to its failure to incorporate the complex reality of socio-ecological systems, the effectiveness of the CITES framework has been questioned in relation to tackling illegal wildlife trade [28, 48]. For instance, the CITES implementation rarely touches grassroot stakeholders (i.e. fishers), who are the most impacted by the regulation and tend to leave them with uncertainty and misinformation. CITES implementation should be periodically evaluated to examine its effectiveness and shifts of behaviour. It is also crucial to investigate any alteration of trade behaviour (i.e. route, volume and source) which may be counter-productive to CITES principles [49]. Without adjustments, coastal communities are unlikely to benefit from CITES implementation, which may instead render their business more uncertain; so a practical alternative is required for communities that depend on CITES species, optimising the benefits while minimizing the costs [50].

Despite the valuable efforts by the B/LPSPL (‘Balai/Loka Pengelolaan Sumber Daya Pesisir dan Laut’; Institute for Coastal and Marine Resource Management) authority of the Ministry for Marine Affairs and Fisheries to meet the three main principles of CITES (i.e. legality, sustainability, and traceability) across the country, limited resources still represent major challenges for authorities and exporters. Species identification is also extremely challenging since sharks and rays are processed in a myriad of ways, which makes the tracing of exports very challenging [26]. All these circumstances determine the intricacy of domestic and international trade flows in Indonesia (**Error! Reference source not found.**), whose disentanglement will require multi-disciplinary approaches, solid collaboration and substantial engagement.

## 5. Conclusion

We have made a major step towards understanding historical and current trends in landing, domestic flow and international trade of sharks and rays in Indonesia. We found that species catch recording, domestic traceability, and international trade are all inadequate to guarantee long-term conservation of these living resources, and also fail to help local communities securing a reliable and sustainable food source. There is also great doubt that the value chain is fair to fishers and local operators, especially concerning the luxury products that are exported. An increase of shark species listed in the CITES Appendices highlights the importance of improving national capability to monitor the supply chain, from net to consumers/importers. The current scenario calls for efforts to be made towards: i) increasing taxonomic resolution of landing and trade statistics, ii) standardisation of product-based HS codes to facilitate consistent naming among authorities (Cawthorn et al 2018); iii) expanding national capabilities in technologies (e.g. DNA testing, Cardeñosa *et al.*, 2018) designed for accurate product identification; iv) taking into account the socio-economic aspects of the fisheries to feed into more effective conservation and management measures.

Community participation is a vital requirement to consider in the early stages of a management plan, and it will also be helpful for the surveillance and stewardship of the management action implemented [51], while conferring the required “ecosystem service elasticity” [52] in the often unique socio-ecological system in question. A typical example is the often touted ‘shark tourism solution’, which is bound to fail without effective community engagement [39].

## Acknowledgements

We thank all collaborators of the project *Building Capacity to reduce illegal trading of shark products in Indonesia*, funded under the Illegal Wildlife Trade Challenge Fund and the University of Salford R&E strategy funding, and based at the Ministry for Marine Affairs and Fisheries (MMAF) – Republic of Indonesia, the University of Salford (UoS), the Centre for Environment, Fisheries and Aquaculture Science (Cefas) and the Wildlife Conservation Society – Indonesian Program (WCS-IP). We also thank to officers of B/LPSPLs (‘Balai/Loka Pengelolaan Sumber Daya Pesisir dan Laut’; Institute for Coastal and Marine Resource Management) and Fish Quarantine for helping during field work.

## Author contribution

A.P., A.D.M., J.M., J.M., F.A, E.M., and S.M. conceived, designed and coordinated the study; Data Collection and data analyses were conducted by A.P; A.P. wrote the manuscript; all authors read and commented on the manuscript.

## Additional Information

### Supplementary information

Supplementary Table S1. Shark and ray production and trade data used in this study

Supplementary Figure S2. Domestic trade network of fin and meat products across Indonesia region within 2014-2018 (ton)

Supplementary Figure S3. Annual volume of reported export and import by/from Indonesia in 2012-2018 for fin products (a) and meat products (b)

Supplementary Table S4: Shark product HS codes used in trade, 2008–2018 (UN Comtrade)

