## Supplement Material for "Disentangling the shark and ray trade in Indonesia to reconcile conservation with food security"

**Supplementary Table S1. Shark and ray production and trade data used in this study**. Trade data include HS Code and descriptions of shark and ray commodities.

| Data source | Information | Designation |
|---|---|---|
| **Production statistics** | | |
| Indonesian Marine and Fisheries in Figure 1975-2016 (MMAF, 2017) | Species, fisheries management area, province, volume | Indonesia classification on sharks and rays |
| One Data of Indonesian fisheries 2017-2018 (MMAF, 2020) | Species, fisheries management area, province, volume | Indonesia classification on sharks and rays |
| FAO Global capture production 1950-2018.<br>Accessed via FishstatJ data (FAO, 2020) | Country, species, volume, value | ISSCAAP group > Sharks, rays, chimaeras |
| **Trade statistics** | | |
| FAO Global Fisheries commodities production and trade 1976-2017.<br>Accessed via FishstatJ data (FAO, 2020) | Flow, source and destination country, commodity, HS code, volume, value | ISSCAAP group > Sharks, rays, chimaeras |

| | | |
|---|---|---|
| Indonesian fish quarantine data 2014-2018.<br>Accessed via online query panels, 2010-2016 (AFQQI-MMAF, 2019) | Flow, source and destination country, commodity, volume, value | Indonesia classification on sharks and rays |

**Supplementary Figure S2. Domestic trade network of fin and meat products across Indonesia region within 2014-2018 (ton)**

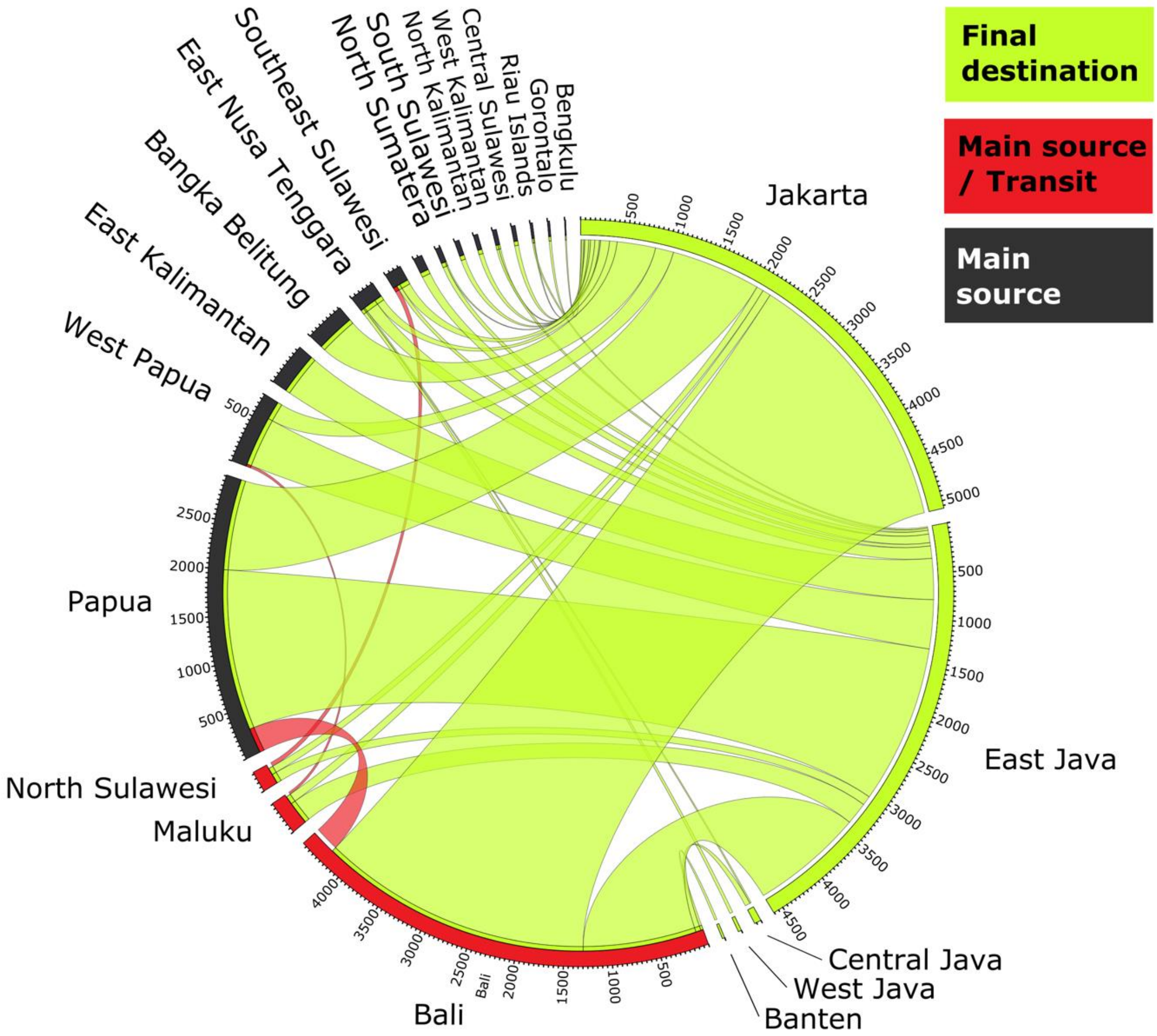

**Supplementary Figure S3. Annual volume of reported export and import by/from Indonesia in 2012-2018 for fin products (a) and meat products (b)**

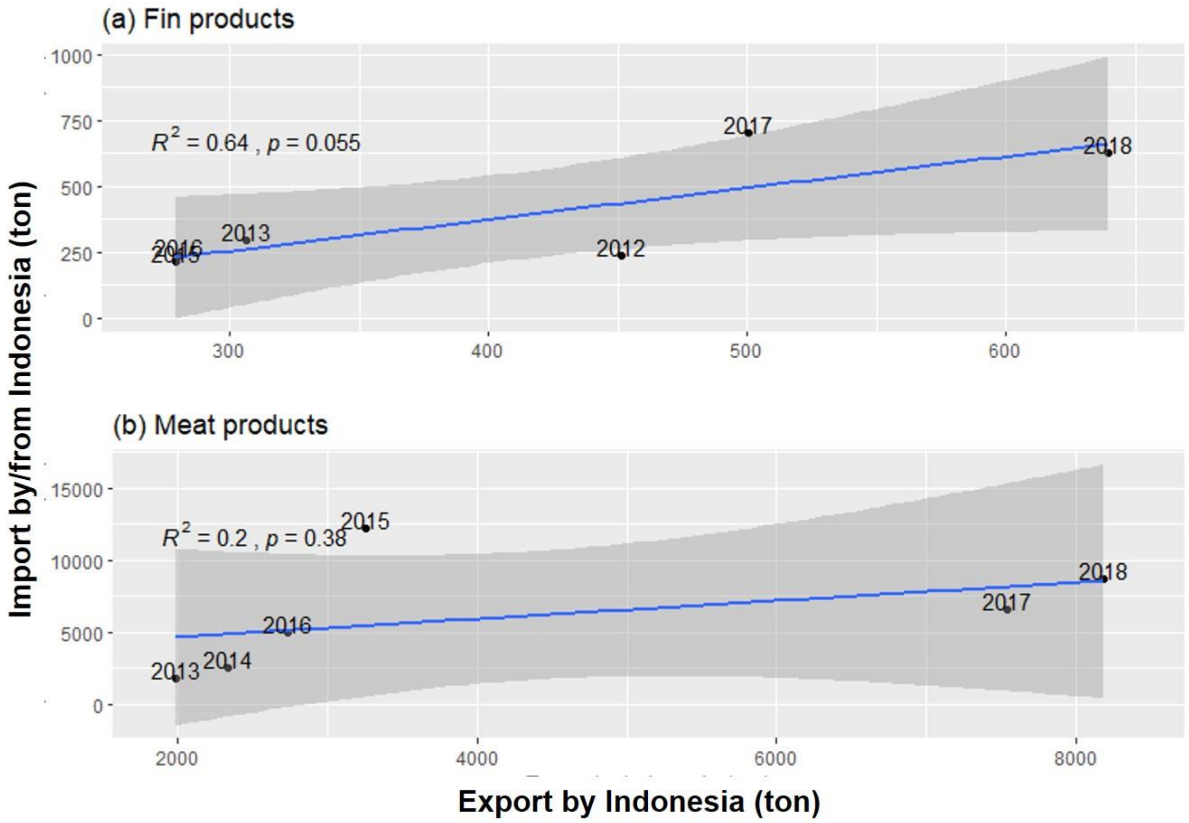

**Supplementary Table S4: Shark product HS codes used in trade, 2008–2018 (UN Comtrade)**

| HS Code | Meat | HS Code | Fins |
|---|---|---|---|
| 03.02.65 | Dogfish & other sharks, fresh/chilled (excl. fillets/other fish meat of 03.04/livers & roes) | 03.02.92 | Fish; fresh or chilled, shark fins |
| 03.02.81 | Fish; fresh or chilled, dogfish and other sharks, excluding fillets, fish meat of 0304, and edible fish offal of subheadings 0302.91 to 0302.99 | 03.03.92 | Fish; frozen, shark fins |
| 03.03.75 | Dogfish & oth. sharks, frozen (excl. fillets/oth. fish meat of 03.04/livers & roes) | 03.05.71 | Fish; edible offal, shark fins |
| 03.03.81 | Fish; frozen, dogfish and other sharks, excluding fillets, fish meat of 0304, and edible fish offal of subheadings 0303.91 to 0303.99 | 1604.18 | Fish preparations; shark fins, prepared or preserved, whole or in pieces (but not minced) |
| 03.04.47 | Fish fillets; fresh or chilled, dogfish and other sharks | | |
| 03.04.56 | Fish meat; excluding fillets, whether or not minced; fresh or chilled, dogfish and other sharks | | |
| 03.04.88 | Fish fillets; frozen, dogfish, other sharks, rays and skates (Rajidae) | | |
| 03.04.96 | Fish meat, excluding fillets, whether or not minced; frozen, dogfish and other sharks | | |

Notes: The Harmonized System (HS) product code is a standardized numerical method of classifying traded products. Those six-digit code (except for 160418) structured into 3 section i.e. chapter (product), heading (type of treatment), and subheading (specify the species). First two-digit stands for fish and crustaceans, molluscs and other aquatic invertebrates. While the next two digits refer to the treatment i.e. 01 if for “live”, 02 is for “fresh or chilled”, 03 is for “frozen”, 04 is for “filleted”, and 05 is for “dried, salted, smoked, and pelleted”. Then, after the first four digits used to specify the species. Meanwhile, 1604 stands for “prepared or preserved fish” and the last two-digit refer to sharks. Additionally, this 6 six-digit international code could be added a national classification code to increase clarity.
